# Ecological characterization and infection of Anophelines (Diptera: Culicidae) of the Atlantic Forest in the southeast of Brazil over a 10 year period: Has the behaviour of the autochthonous malaria vector changed?

**DOI:** 10.1101/146803

**Authors:** Julyana Cerqueira Buery, Helder Ricas Rezende, Licia Natal, Leonardo Santana da Silva, Regiane Maria Tironi de Menezes, Blima Fux, Rosely dos Santos Malafronte, Aloisio Falqueto, Crispim Cerutti

## Abstract

In the south and southeast of Brazil, autochthonous malaria cases can be found near Atlantic Forest fragments. The transmission is not totally clarified; thus, the behaviour of the possible vectors in those regions must be observed. An entomological and natural infection study was performed on anophelines (Diptera: Culicidae) captured in the municipalities of the mountainous region of Espírito Santo state in 2004-2005. Similarly, between the years 2014 and 2015, 12 monthly collections were performed at the permanent trapping station of the study mentioned above (Valsugana Velha, Santa Teresa, ES). Light traps with CO_2_ (CO_2_-baited Center for Disease Control [CDC] traps) were set in open areas, at the edge of the forest (canopy and ground) and inside the forest (canopy and ground), whereas Shannon traps were set on the edge of the forest. A total of 1,414 anophelines were collected from 13 species. *Anopheles (Kerteszia) cruzii* Dyar and Knab remained the most captured species in the CO_2_-baited CDC traps set in the forest canopy and was also the vector with the highest prevalence of *Plasmodium vivax* infection according to molecular PCR techniques. Regarding mosquitoes of the subgenus *Nyssorhynchus*, *P. vivax* was found only in abdomens, weakening the hypothesis that this subgenus also plays a role in malaria transmission in this specific region.

**Sponsorship:** Espírito Santo Research Foundation (Fundação de Amparo à Pesquisa e Inovação do Espírito Santo – FAPES).

## INTRODUCTION

Despite being a highly prevalent disease in the Amazon region, malaria remains residual in Atlantic Forests of the south and southeast regions of Brazil. In this region, the disease is known as bromeliad-malaria since the vectors of the genus *Anopheles* reproduce in the whorls of *Bromeliaceae*, which are typical plants of this biome (Downs & Pittendrigh, 1946). The presence of a few autochthonous (sometimes asymptomatic) human cases, the spatial distance between the cases and the low parasitemia by microscopy weaken the possibility of the traditional transmission chain. It is believed that the cycle of *Plasmodium spp.* is maintained by reservoirs of the parasite, in the forest or in the rural population, represented by both apes and humans, asymptomatic or not. Studies have been conducted to clarify these beliefs (Curado et al., 1997; Duarte et al., 2006; Cerutti et al., 2007; Rezende et al., 2009; Meneguzzi et al., 2009). Regarding the vector, in the state of Espírito Santo, Brazil, 26 species of *Anopheles sp.* have been identified, indicating an abundance of anophelines (Coutinho, 1947; Andrade& Brandão, 1957; Ferreira et al., 1964; Natal, 2007; Sallum, 2008; Rezende et al. 2009; Meneguzzi et al. 2009; Silva et al., 2013). The vectors of greater vectorial capacity and competence belong to the subgenus *Kerteszia*, mainly *Anopheles cruzii*, in the south and southeast Brazilian states (Meneguzzi et al., 2009). However, species of anophelines involved in the dynamics of malaria transmission outside the Amazon region vary according to environmental and epidemiological conditions (Pina-Costa et al., 2014). The autochthonous cases of extra-Amazonian malaria are found in mild mountainous regions and are associated with agricultural activities near the forest that are performed by young men (Cerutti Jr et al., 2007). States such as São Paulo, Santa Catarina and Espírito Santo are covered by dense Atlantic Forest regions (IESB, 2007). Among them, Espírito Santo has recorded the largest number of bromeliad-malaria cases in recent years. The biome in question is very humid, with abundant rainfall and vegetation that favours the reproduction of the anophelines. In addition, bromeliad-malaria is beginning to be used as a biological marker of human activities within these forests (Gomes, 1985; Rezende et al., 2013). Rezende et al. (2009) found anopheline behaviour similar to that expected, based on the literature, in Valsugana Velha, Santa Teresa municipality: a high prevalence of *A. cruzii* was reported in a mountainous region of the Atlantic Forest, in an endemic area of the disease in the state of Espírito Santo. However, considering the hypothesis of a typical transmission model of zoonosis in this scenario, this model would require a stable and constant vectorial behaviour. Such behaviour, therefore, must be re-evaluated to establish transmission characteristics over the years.

## MATERIALS AND METHODS

### Description of the study site

The study was performed in the rural area of the municipality of Santa Teresa, which is located approximately 78km from Vitória, in the state of Espírito Santo (ES), Brazil. The permanent trapping station is in Valsugana Velha (19°57′58.4 “S, 40°34′45.2” W), where cases of residual malaria from Atlantic Forest systems were recorded in the study by Cerutti Jr. et al. (2007) (Figure 1).

**Figure 1.**
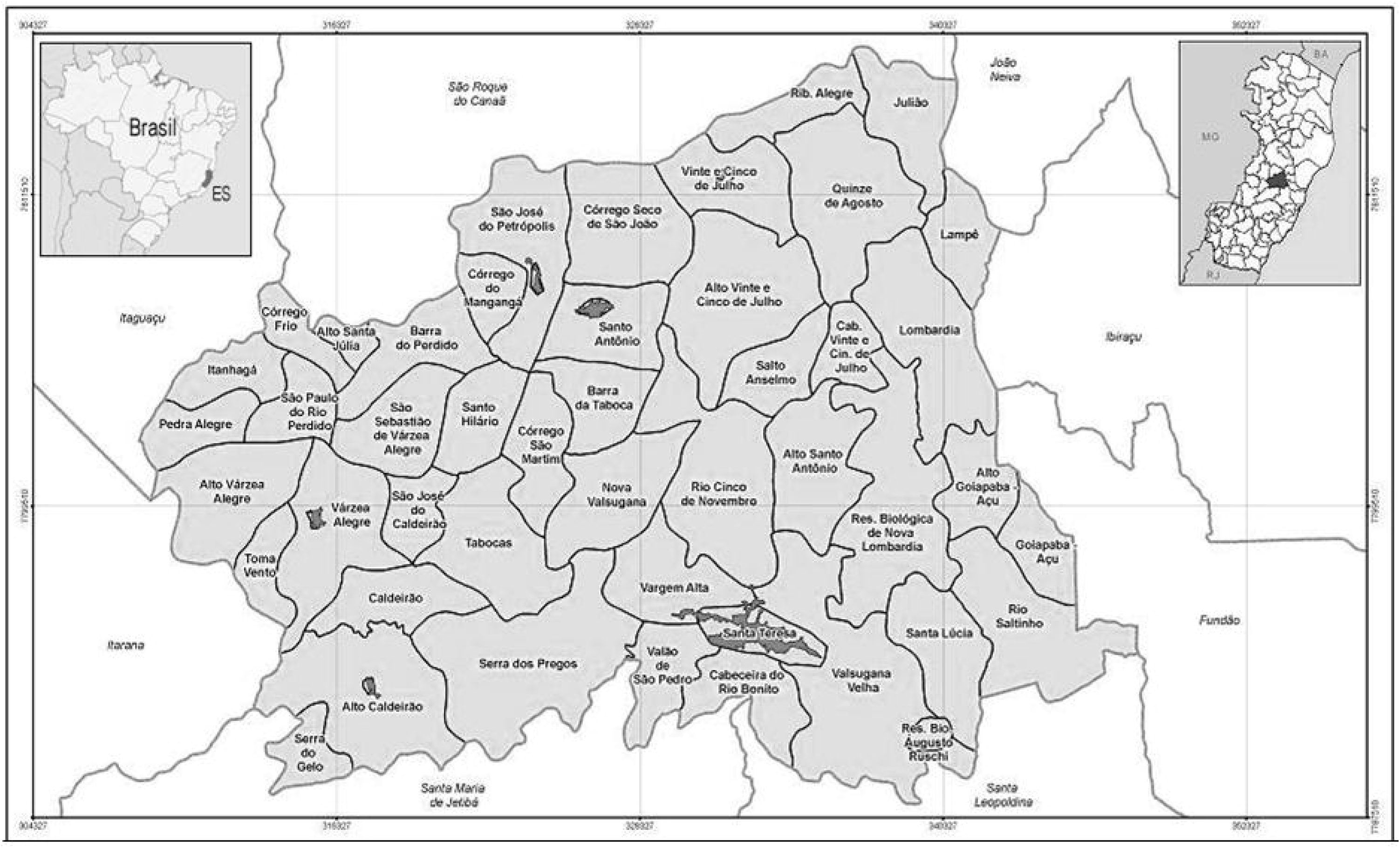
Political map of the municipality of Santa Teresa, highlighting the rural community where the collections were performed. Source: Instituto Jones dos Santos Neves, 2013.

[Figure 1. Political map of the municipality of Santa Teresa, highlighting the rural community where the collections were performed. Source: Instituto Jones dos Santos Neves, 2013.]

### Collection of anophelines

Hourly collections of anophelines were conducted at a permanent trapping station located in the bromeliad-malaria transmission area. These collections were performed one day each month for one year, from June 2014 to May 2015, totalling 12 collections. Two capture methods were used:

- Center for Disease Control (CDC) light traps with CO_2_ (Gomes, 1985) that were set in open areas (peridomiciliary environment = PD), at the edge of the forest (canopy and ground) and inside the forest (canopy and ground) and
- Shannon traps (Bustamante, 1951), set at the edge of the forest.

The five CDC traps were set simultaneously, two of them at 15 metres from the ground in the canopy (on the edge and inside the forest), two at one metre from the ground (at the edge and inside the forest), and one at the border between the forest and the area close to the dwellings. The collections lasted for 12 hours, with the traps placed at night (6:00 PM) and removed in the morning (6:00 AM). For the Shannon traps, the insects were captured during the first 4 hours after sunset (6:00 PM to 10:00 PM) each month.

### Storage and identification of insects

The specimens were stored in tubes containing isopropanol and later identified using the identification keys proposed by Consoli&Lourenço-de-Oliveira (1994). The identification was made by a team from the Entomology and Malacology Centre of Espírito Santo (Núcleo de Entomologia e Malacologia do Espírito Santo - NEMES/ES).

### Molecular techniques

The DNA for the detection of *Plasmodium sp.* was obtained from the thorax, abdomen or entire mosquito of the mosquito groupings in pools (maximum of 10 samples/pool), depending on the subgenus of the specimens. Those of the subgenus *Nyssorhynchus* were sectioned, whereas those of the subgenus *Kerteszia* were processed in whole. The same extraction kit (DNeasy Blood & Tissue Kit, Qiagen, Germany) was used, following the manufacturer’s instructions. Each pool included females of the same species, collected in the same type of trap, on the same date. The presence of *P. vivax* or *P. malariae* in the pools was detected using the nested-PCR protocol described by Kimura et al. (1997) and modified by Win et al. (2002). The target was the gene encoding the 18S ribosomal subunit. The amplification products were analysed by electrophoresis in 2% agarose gel under ultraviolet light. In positive cases, 100-bp fragments were amplified.

### Statistical analysis

To determine the importance and distribution of the various *Anopheles* species, analyses of diversity, dominance and abundance were performed using Shannon’s diversity index (H′) and Simpson’s dominance index (D). In the various comparisons, the level of significance was set at 5%. Bivariate Spearman correlation calculations were used to determine the relationship between anopheline capture, temperature and rainfall (data provided by the Capixaba Institute of Research, Technical Assistance and Rural Extension [Instituto Capixaba de Pesquisa, Assistência Técnica e Extensão Rural - INCAPER]), also at a level of 5%.

## ETHICAL STANDARDS

During the collection process, no harm was inflicted to the environment. The team members wore long clothing, gloves and hats with head nets to avoid anopheline mosquito bites. Boots were also used to prevent bites by venomous animals. A license to capture arthropod insects was obtained via the SISBIO/Chico Mendes Institute (ICMBio/IBAMA/MMA) under number 19227-1.

## RESULTS

A total of 1,414 specimens were captured, resulting in a set of 13 species. The largest number of specimens was collected in April 2015 (341) and the smallest number of specimens in March 2015 (05). In all collections, there was a predominance of *Anopheles (K.) cruzii*, totalling 1,044 of the 1,414 mosquitoes captured (Table 1). The trap in which most mosquitoes were captured was the CO_2_-baited CDC trap placed in the canopy, in which *A. cruzii* was also predominant (Graph 1).

**Table 1.**
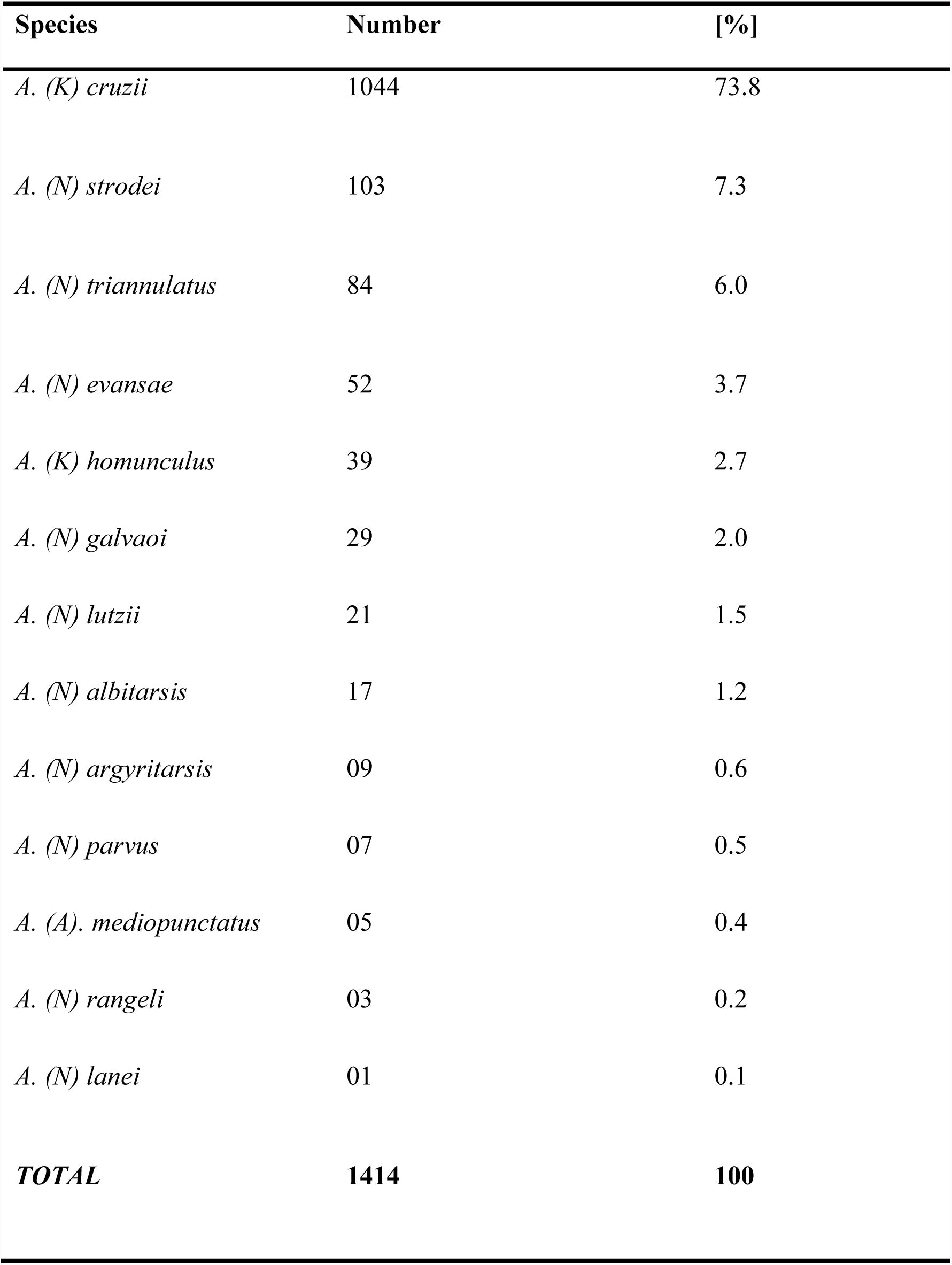
Percentage of anopheline species found between June 2014 and May 2015 at the permanent trapping station in Santa Teresa, ES.

[Graph 1. Percentage of species captured per trap, from June 2014 to May 2015, at the permanent trapping station in Santa Teresa, ES.]

### Climate

The months of November 2014, September 2014 and April 2015 showed the highest capture frequencies (Graph 2). There was no simultaneous capture of all species in the monthly collections, and all species were absent for at least 1 month during the total collection period. *A. cruzii* was not captured only in June 2014 and March 2015 (Graph 2). In fact, those were the only months in which *A. cruzii* ceased to be the predominant species and was replaced with *A. strodei* and *A. triannulatus*, respectively.

[Graph 2. Species captured each collection month, from June 2014 to May 2015, at the permanent trapping station of Santa Teresa, ES.]

A negative trend was observed for the correlation between the number of anophelines of the subgenus *Nyssorhynchus* and the temperature, although without statistical significance (r = - 0.04, p = 0.89). Additionally, a positive trend was observed for the correlation between the number of anophelines and rainfall (r = 0.17, p = 0.59) but also with no statistical significance. For the subgenus *Kerteszia*, the correlation between capture frequency and temperature (r = -0.004, p = 0.99) or rainfall (r = -0.13, p = 0.68) revealed negative coefficients but with no statistical significance. The monthly average temperature and the rainfall in Valsugana Velha are shown in Graph 3.

[Graph 3. Temperature and rainfall data in Santa Teresa, ES, between June 2014 and May 2015.]

### Spatial distribution

*A. strodei* and *A. triannulatus* were more frequently captured in the Shannon traps, and *A. cruzii* was captured more frequently in the CO_2_-baited CDC traps (Graph 1) located in the tree canopy. Simpson’s dominance index (D) reveals that dominance in the Shannon trap (D1 = 0.227) was greater than the dominance of individuals collected in the CO_2_-baited CDC trap (D2 = 0.172) (p < 0.02), both at the forest edge.

Based on Shannon’s diversity index (H′), the diversity of anophelines collected in the Shannon trap (H′2 = 1.866) was higher compared to that of the anophelines collected in the CDC trap at the forest edge, near the dwellings (H′1 = 1.734) (p = 0.004). *Anopheles strodei* was captured in larger numbers at forest edge areas, followed by *A. triannulatus. A. lanei* and *A. rangeli* were captured only in the CO_2_-baited CDC trap located near the dwellings, whereas *A. homunculus* was only captured in the CDC traps set in the tree canopy.

### Natural infection rates

(Table 2). Of the total pool of specimens investigated, 13 were positive for *Plasmodium vivax* (Figure 2). Of these, 10 belonged to the species *A. cruzii*, and three were abdomens that belonged to specimens of the subgenus *Nyssorhynchus*. As shown in Table 2, most of the infected mosquitoes were *A. (K.) cruzii*, captured in CO2-baited CDC traps located in the tree canopy. Graph 4 shows the monthly distribution of captured mosquitoes in which *P. vivax* was found. *P. malariae* infection was not detected.

**Table 2.** Species, traps and collection dates for *P. vivax-*positive mosquitoes at the permanent trapping season in Santa Teresa, ES.

| Sample | Species | Trap | Date |
| --- | --- | --- | --- |
| **260a\*** | *A. (N.) lutzii* | Shannon | September 2014 |
| **279** | *A. (K.) cruzii* | CDC canopy | August 2014 |
| **343** | *A. (K.) cruzii* | CDC canopy | August 2014 |
| **479** | *A. (K.) cruzii* | CDC canopy | October 2014 |
| **485** | *A. (K.) cruzii* | CDC canopy | October 2014 |
| **494a\*** | *A. (N.) evansae* | Shannon | October 2014 |
| **632** | *A. (K.) cruzii* | CDC ground | November 2014 |
| **633** | *A. (K.) cruzii* | CDC canopy | November 2014 |
| **868** | *A. (K.) cruzii* | CDC canopy | February 2015 |
| **1142** | *A. (K.) cruzii* | CDC canopy | April 2015 |
| **1144** | *A. (K.) cruzii* | CDC canopy | April 2015 |
| **1248** | *A. (K.) cruzii* | CDC canopy | April 2015 |
| **1294a\*** | *A. (N.) strodei* | CDC near dwellings | May 2015 |

\*a = abdomen.

**Figure 2.**
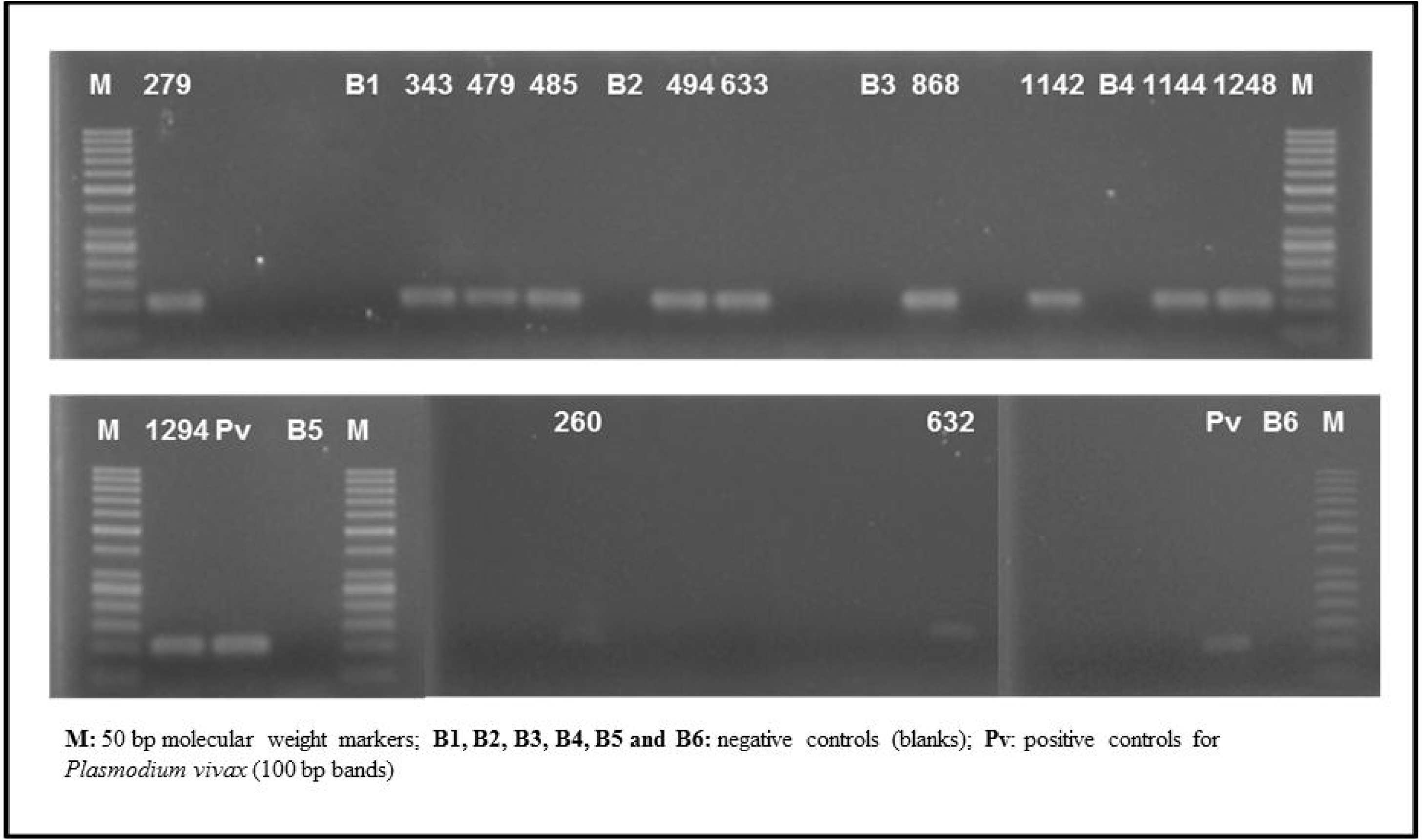
Ethidium bromide-stained agarose gel, photographed using the Alpha Imager software, with 13 samples of pools of *P. vivax*-positive anopheline females. Source: Buery JC and Natal L, 2016

[Figure 2. Ethidium bromide-stained agarose gel, photographed using the Alpha Imager software, with 13 samples of pools of *P. vivax*-positive anopheline females. Source: Buery JC and Natal L, 2016.]

[Graph 4. Chronological distribution of *P. vivax*-infected anopheline females captured in the permanent trapping station in Santa Teresa, ES.]

## DISCUSSION

Bromeliad malaria became more evident in the Brazilian south and southeast Atlantic forest regions after control measures implemented by the government eliminated transmission in flat areas exploited by humans (Ministério da Saúde (BR), 2015; Barata, 1998). As residual foci remained in the eradication areas, bromeliad-malaria or malaria of Atlantic Forest systems began to be studied with more intensity in the 1980s and 1990s (Duarte et al., 2013). With a more consistent understanding of vector ecology and outbreak epidemiology, more evidence has emerged in favour of behaviour consistent with a zoonosis. The zoonosis hypothesis presupposes the concomitant existence of simian malaria. Such an existence has been proven in the past (Duarte et al., 2008), but its prospective evaluation demands extremely complex logistics. Therefore, the evaluation of vector behaviour favourable to enzootic transmission is more feasible and may provide indirect evidence of the presence of the infection among monkeys and of its source of occurrence in human cases. Given the need for stable and constant vector behaviour for the maintenance of simian malaria, follow-up studies of the endemic regions are necessary. In addition, according to Marrelli et al. (2007), the entire Atlantic Forest territory that survived deforestation must be carefully monitored due to environmental changes and the possibility of maintaining the infection cycle given the high number of asymptomatic individuals who can act as reservoirs of the parasite in those regions.

In the present study, *A. cruzii* still prevailed as the main vector found at the Valsugana Velha trapping station. Between 2004 and 2005 (Rezende et al. 2009), 61.2% of the 2,290 mosquitoes collected belonged to this species. In 2014 and 2015, 73.8% of the 1,414 anophelines captured were *A. cruzii*. Therefore, an increase in the proportion of the main vector of bromeliad malaria was observed in the local fauna. Older studies such as those of Deane (1988) and more recent studies such as those of Duarte et al. (2013), Neves et al. (2013) and Kirchgatter et al. (2014) have previously demonstrated the magnitude of the presence of this species in Atlantic Forest regions with preserved native forest.

The distribution of *A. cruzii* when moving from the inside to the edge of the forest has also remained the same over time. As Rezende et al. (2009) described in 2004 and 2005, the anophelines of this species appeared mostly in the tree canopy inside the forest, whereas the numbers decreased drastically in the traps that were set closer to the forest edge and the dwellings. The anopheline fauna was dominated by species such as *A. strodei* and *A. triannulatus* closer to human-occupied areas. In São Paulo, between 2009 and 2011, this trend was also detected when comparing anthropic and wild areas in the town of Parelheiros. There, 438 *A. cruzii* were recorded in a certain anthropic area and 4,832 in the wild area (Duarte et al., 2013).

The absolute predominance of *A. cruzii* in the canopy, observed in the present study, indicates a non-synanthropic behaviour (Forattini et al., 1990) and reinforces its role in the transmission of simian malaria. However, despite being present mostly in the canopy, some specimens were found in the CO_2_-baited CDC traps set in the ground and in the Shannon traps, which points to the possibility of these vectors descending from their preferential site in the canopy, at which time they could incidentally transmit the parasites to humans. The presence of these specimens reinforces the hypothesis that bromeliad-malaria is maintained in the region by means of simian infection. The fact that humans are in the forest and that *A. cruzii* comes down from the canopy to feed creates the conditions for *Plasmodium spp.* originating from the monkeys in the canopy to infect humans.

The morphometric diagnosis studies of Lorenz et al. (2012) showed a differentiation between *A. cruzii* and *A. homunculus* not predicted in the study of Rezende et al. (2009). However, only 39 *A. homunculus* were captured during the collection period between 2014 and 2015, against 1,045 duly identified *A. cruzii*.

Unexpectedly, the season during which most mosquitoes were captured was not the summer. Interestingly, the seasons with milder weather had the largest capture rates. For example, the most successful captures occurred in the months of September 2014 and April 2015. This result corroborates the study of Rezende et al. (2009), who suggested the adaptation of anophelines to mild environments. There was also a high number of specimens collected in November 2014. The mean rainfall at that time was the highest of that year (mean of 207.6 mm/month), and the temperature was mild on collection day (ranging from 19.7 to 21.3°C). That month, the “rain” factor may have triggered an increase in the mosquito population.

In this study, 13 pools of mosquitoes were positive for *P. vivax*. In 2004 and 2005, 10 pools of anophelines infected by the same species of the parasite were obtained (Rezende et al., 2009). However, unlike 10 years ago, the PCR reactions for the 2014 and 2015 collections did not show infection in the thorax of mosquitoes of the subgenus *Nyssorhynchus*. The infective form should reach the salivary gland of the anopheline for infection to occur. Thus, the separation into thorax and abdomen during the experiments reinforces evidence of the participation of other vector species in the transmission chain. Because *Kerteszia* was a subgenus of a known vector, there was no separation of the body to perform the experiments, as occurred in mosquitoes of the subgenus *Nyssorhynchus*, to assess the possibility that these mosquitoes participate in the transmission chain. Since no infection was detected in the *Nyssorhynchus* thoraxes, unlike 10 years ago, *Nyssorhynchus* may have stopped playing the role of secondary vector and *A. cruzii* may currently be the only bromeliad-malaria vector in this area. The progressive exploitation of the rural and forest environment by the local inhabitants may have led to greater spatial distances between the transmission events and the anthropic environment, where *Nyssorhynchus* has greater dominance. In these conditions, given the *A. cruzii* dominance, *Nyssorhynchus* does not have the opportunity to become infected. Humans venture into the forest to clean the river springs or to gather firewood and thus acquire the disease. Once they return to their homes, they likely infect the *Nyssorhynchus* near their dwellings.

Infected *A. cruzii* were collected in CO_2_-baited CDC traps in the tree canopy inside the forest and in one CO_2_-baited CDC trap located near the ground, at a height of one metre. This fact reinforces the idea that infections can occur in both habitats as a result of the acrodendrophilic behaviour with eventual descent to lower heights, where they can feed on non-usual hosts. This fact also reinforces the possibility of the disease being a zoonosis in regions such as Valsugana Velha, in Santa Teresa, ES.

Regarding the Shannon’s dominance index (H′), there was greater dominance in the Shannon trap than in the CO_2_-baited CDC trap, both at the forest edge.

Regarding Simpson’s diversity index (D), the diversity of anophelines collected with the Shannon trap was also higher than that observed in the CO_2_-baited CDC trap in the same habitat. The placement of the trap near the body of water, where there were *Nyssorhynchus* breeding sites, would justify the greater diversity.

These data corroborate a study conducted in the same region and published in 2013 (Rezende et al., 2013), when greater dominance and diversity of anophelines were observed in an anthropic environment with the presence of malaria. These findings reinforce the role of human occupation in the determination of both anopheline distribution and behaviour since both indices show higher values in collections performed in an environment closer to human dwellings.

The study shows that there was little change in vector behaviour in the region studied. *A. cruzii* remains the most infected anopheline in Valsugana Velha, and mosquitoes of the subgenus *Nyssorhynchus* do not appear to participate in the transmission chain. The acrodendrophilic behaviour of *A. cruzii*, particularly those infected, reinforces the hypothesis that the presence of *P. vivax* in these specimens arises from blood feeding in animals that live in the tree canopy, such as simians.

## ACKNOWLEDGEMENTS

We would like to thank the State Health Department of Espírito Santo and the Entomology and Malacology Centre of Espírito Santo/SESA for the logistical support and supply of equipment in the field; Claudiney Biral dos Santos for technical support; Filomena E C Aguiar, Creuza Rachel Vicente, Marcelo Urbano Ferreira, Priscila Thihara Rodrigues and Lais Camoese Salla for the critical reading of the manuscript; the owners of the Recanto da Preguiça farm for allowing sampling at that site; and the Capixaba Institute of Research, Technical Assistance, and Rural Extension

(INCAPER/ES) for providing the rainfall and temperature data.

---

**Graph 1.**
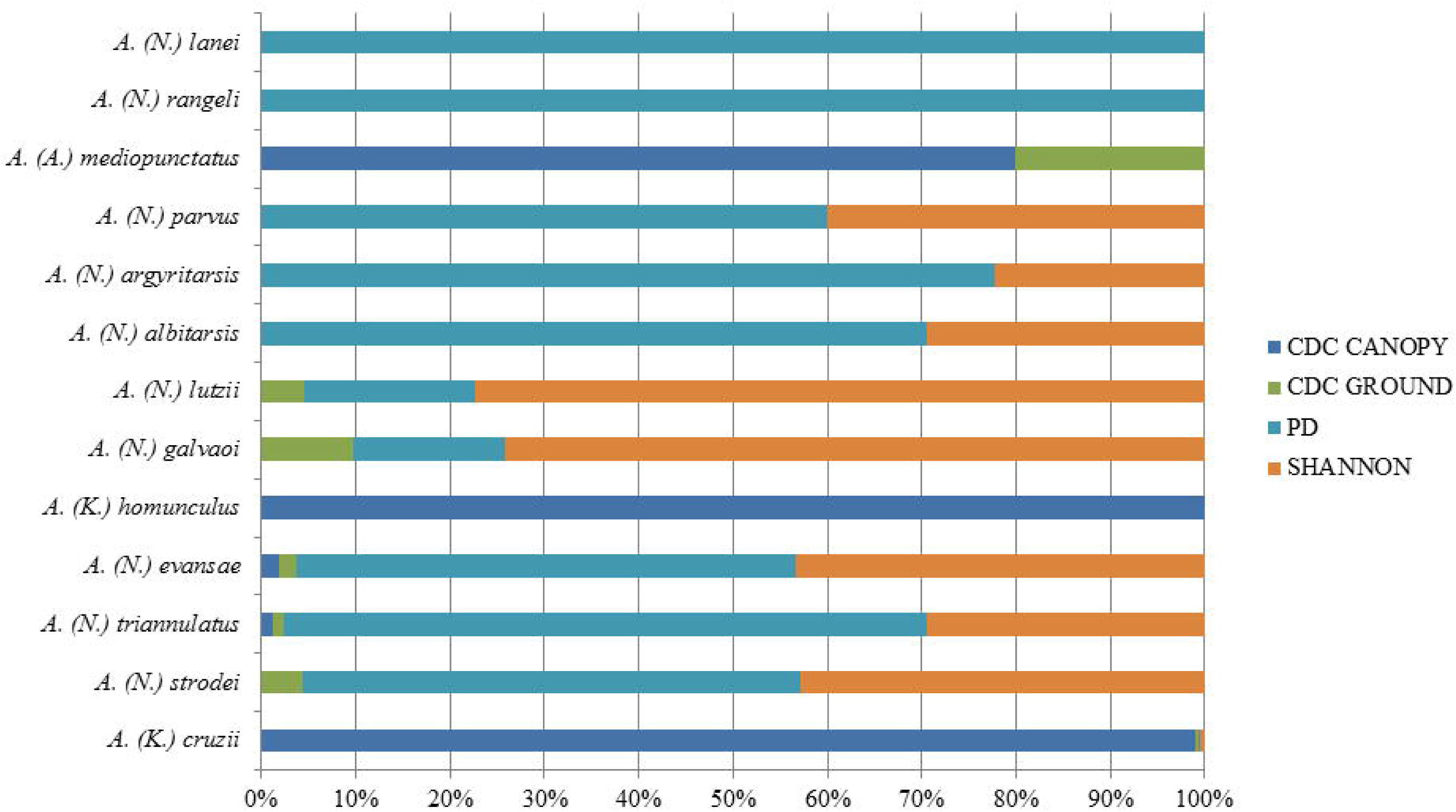
Percentage of species captured per trap, from June 2014 to May 2015, at the permanent trapping station in Santa Teresa, ES.

**Graph 2.**
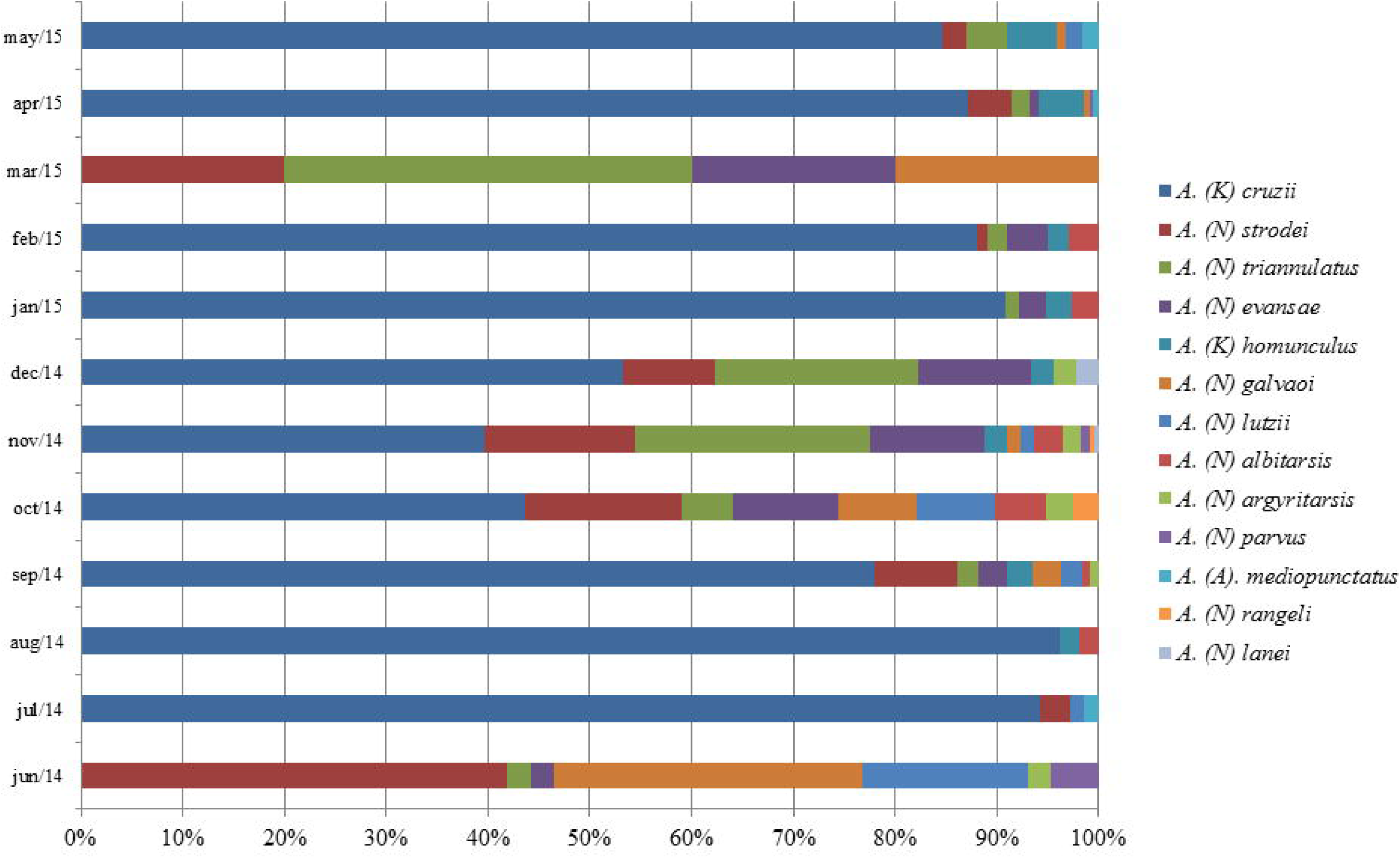
Species captured each collection month, from June 2014 to May 2015, at the permanent trapping station of Santa Teresa, ES.

**Graph 3.**
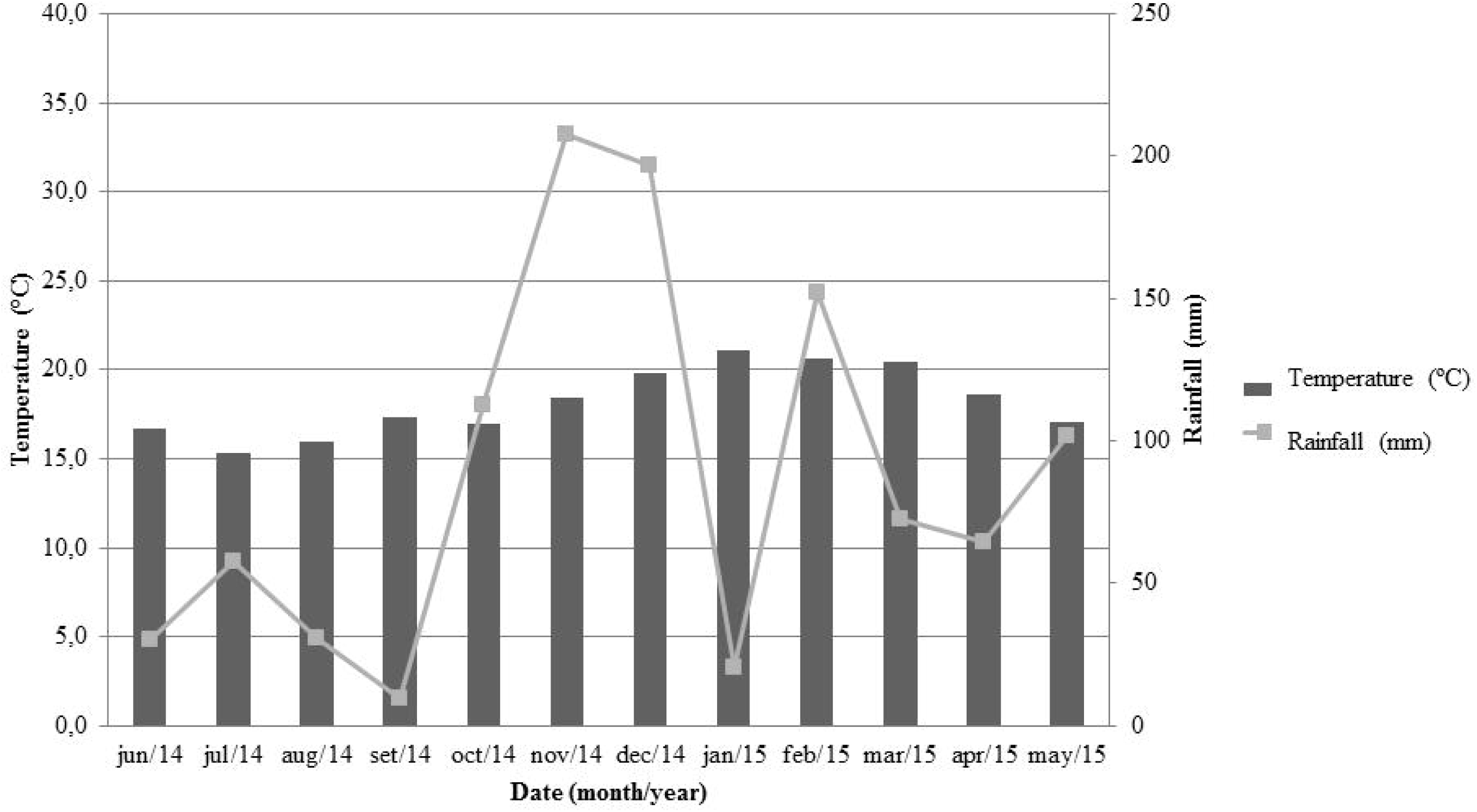
Temperature and rainfall data in Santa Teresa, ES, between June 2014 and May 2015.

**Graph 4.**
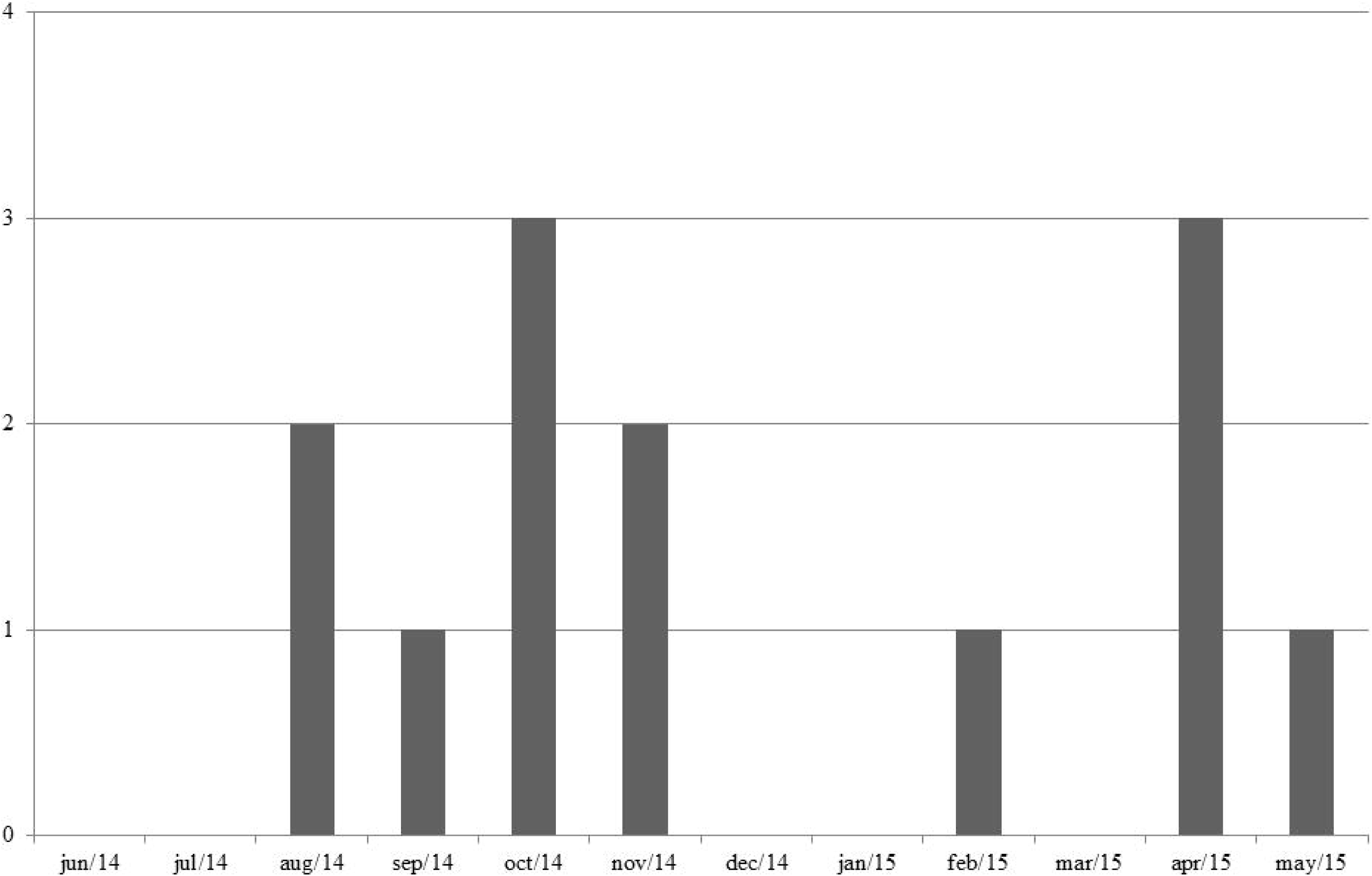
Chronological distribution of *P. vivax*-infected anopheline females captured in the permanent trapping station in Santa Teresa, ES.

